# Magnetically-Propelled Fecal Surrogates for Modeling the Impact of Solid-Induced Shear Forces on Primary Colonic Epithelial Cells

**DOI:** 10.1101/2021.02.18.431336

**Authors:** Samuel S. Hinman, Jennifer Huling, Yuli Wang, Hao Wang, Ross C. Bretherton, Cole A. DeForest, Nancy L. Allbritton

**Affiliations:** Department of Bioengineering, University of Washington, Seattle, WA, USA; Joint Department of Biomedical Engineering, University of North Carolina at Chapel Hill, Chapel Hill, NC, USA, and North Carolina State University, Raleigh, NC, USA; Department of Chemical Engineering, University of Washington, Seattle, WA, USA

**Keywords:** colon, intestine, primary cells, shear force, magnetic, hydrogel

## Abstract

The colonic epithelium is continuously exposed to an array of biological and mechanical stimuli as its luminal contents are guided over the epithelial surface through regulated smooth muscle contraction. In this report, the propulsion of solid fecal contents over the colonic epithelium is recapitulated through noninvasive actuation of magnetic agarose hydrogels over primary intestinal epithelial cultures, in contrast to the vast majority of platforms that apply shear forces through liquid microflow. Software-controlled magnetic stepper motors enable experimental control over the frequency and velocity of these events to match *in vivo* propulsive contractions, while the integration of standardized well plate spacing facilitates rapid integration into existing assay pipelines. The application of these solid-induced shear forces did not deleteriously affect cell monolayer surface coverage, viability, or transepithelial electrical resistance unless the device parameters were raised to a 50× greater contraction frequency and 4× greater fecal velocity than those observed in healthy humans. At a frequency and velocity that is consistent with average human colonic motility, differentiation of the epithelial cells into absorptive and goblet cell phenotypes was not affected. Protein secretion was modulated with a two-fold increase in luminal mucin-2 secretion and a significant reduction in basal interleukin-8 secretion. F-actin, zonula occludens-1, and E-cadherin were each present in their proper basolateral locations, similar to those of static control cultures. While cellular height was unaffected by magnetic agarose propulsion, several alterations in lateral morphology were observed including decreased circularity and compactness, and an increase in major axis length, which align with surface epithelial cell morphologies observed *in vivo* and may represent early markers of luminal exfoliation. This platform will be of widespread utility for the investigation of fecal propulsive forces on intestinal physiology, shedding light on how the colonic epithelium responds to mechanical cues.

**HIGHLIGHTS:**

- Magnetic nanoparticle-embedded agarose hydrogels were developed as surrogates for human feces
- Software-controlled magnetic motors enable programmable studies of colonic motility
- Propulsive shear forces were optimized for primary cell viability, surface coverage, and electrical resistance
- Mucus production and cytokine secretion were modulated by magnetic agarose propulsion
- Structural and cytoskeletal proteins remain properly distributed with alterations in lateral cell morphology

## 1. Introduction

The colon plays several key roles in human health, influencing nutritional uptake, waste management, salt and water homeostasis, drug metabolism, and microbiome composition [1], all of which are supported by the intestinal epithelium [2, 3]. Mechanical forces within the colon play a role in homeostatic and protective mechanisms, including those involved in preventing colorectal cancer, inflammatory bowel diseases, and bacterial infection [4]. Changes in mechanical cues from the local environment have been shown to cause fibrogenic response of colonic fibroblasts [5] and increased proliferation of colonic epithelial stem cells through yes-associated protein 1 (YAP)-dependent mechanisms [6]. On a larger scale, the epithelium experiences smooth muscle contractions that generate intraluminal pressure variations and shear stresses as luminal contents in the form of chyme or feces are guided over the epithelial surface [7, 8]. These organ-level mechanical forces have been suggested to affect serotonin release and intestinal fluid secretion [9, 10] though the exact mechanisms underlying their effects remain unclear due to limited experimental models.

Microphysiological systems generated from primary cells are ideal for studies of organ-level mechanical forces in that they are predictive of human physiology, are highly scalable, and can provide exquisite experimental control that is unavailable in animal models [11]. Several features of *in vivo* colonic motility must be recapitulated, however, in order to accurately mimic normal physiology. Intermittent smooth muscular contractions in the colon trigger fecal movement via peristaltic waves, which are governed by circadian rhythm and eating/fasting schedules [3, 12]. Low-amplitude propagated contractions (wave amplitude *ca*. 20 mm Hg) occur at a constant rate of about 60 contractions d^-1^, assisting in localized mixing and transport of liquids or gas, though not significantly contributing to the bulk movement of luminal contents [12, 13]. High-amplitude propagated contractions (wave amplitude *ca*. 100 mm Hg), on the other hand, occur at a less frequent rate of 4-6 contractions d^-1^ in healthy humans and are associated with colonic emptying, propelling fecal material with a linear velocity of 10 mm s^-1^ [7, 12, 14]. Very little data exists regarding the rheological properties of human feces or the shear stresses imparted by fecal movement, with order of magnitude shear stress estimates ranging anywhere from 10^−3^-10^3^ Pa [15-18]. Additionally, fecal viscosity and associated shear stresses are known to be highly dependent on water content [16-18], which will vary between individuals depending on their diets and disease states [19]. Based on the above *in vivo* benchmarks, the ideal model for investigating the effects of fecal shear forces should exhibit zero to low luminal flow rates at most times, except during the infrequent periods in which solid material is guided over the epithelial surface. While the mechanical properties, propulsion frequency, and linear velocity of model fecal surrogates should match those observed in healthy humans, the ability to experimentally modulate these factors would enable mechanistic studies of pathological conditions in which high-amplitude propagated contractions are adversely affected, as is the case for slow-transit constipation [14] and chronic diarrhea [20].

Several technologies have been developed to examine the various mechanical forces that are present within the small and large intestines. The actuation of electro-responsive membranes has been used to mimic the longitudinal and circular constriction of smooth muscle around the small intestine [21]. Microfluidic organ-on-chips and Flexcell bioreactors are capable of inducing cyclic strain to monolayers of cells cultured on a flexible support, thus also replicating the rhythmic peristalsis of the small intestine [22-26]. To generate luminal shear forces, many systems employ fluid microflow over the epithelial surface, imparting shear stresses of 0.2 – 8.0 mPa [22, 23, 27-29]. However, unlike the small intestine, the luminal contents of the colon are comparatively solid and static due to the organ’s roles in water absorption and waste management. To the best of our knowledge, only one platform has attempted mechanical stimulation of colonic epithelial cells using a solid material, actuating glass microelectrodes against individual enterochromaffin cells to study a mechanosensitive ion channel protein [10, 25]. As such, the impact of solid-induced shear stresses at velocities and frequencies that match those of healthy human fecal contents remain entirely uncharacterized.

Several advances in primary cell culture techniques enable facile *in vitro* expansion of primary adult intestinal stem cells [30-32]. Unlike tumor-derived cell lines, primary intestinal stem cells carry a normal genotype and are capable of generating all differentiated cell lineages that are present *in vivo*, and are thus more predictive of healthy human physiology [33]. Once extracted from primary tissue, long-lived proliferative cells can be subjected to serial expansion over collagen hydrogel scaffolds or within Matrigel patties, maintaining a stable karyotype while in culture [32, 34]. When cultured over porous membranes, these primary epithelial cells form a high-resistance barrier as occurs *in vivo* and can be used for high-content analytical screens in which the basal and luminal compartments are segregated [35-37]. In this report, this primary cell-derived culture format is extended to include the application of solid-induced shear forces, rather than those induced by liquid flow, at user-defined velocities and frequencies that are representative of human colonic luminal contents. Magnetic hydrogels actuated by software-controlled magnetic stepper motors were incorporated as programmable analogs for human feces, enabling experimental manipulation of the velocity and speed of shear force events. These parameters were optimized for primary intestinal epithelial cell culture based on observed alterations in cell surface coverage, viability, and barrier integrity. Parameters matching high-amplitude propagated contractions of the human colon were selected for further analysis, investigating their effects on differentiated lineage allocation, protein secretion (*i*.*e*., mucin-2, interleukin-8), and changes in cell morphology compared to cells grown without shear force exposure. The main strengths of this platform include its compatibility with standard laboratory equipment (*e*.*g*., plate readers, automated microscopes, liquid handling systems) and primary intestinal epithelial cell culture, ensuring that any assays developed will be scalable, reproducible, and predictive [11, 38].

## 2. Materials and methods

### 2.1. Magnetic motorized culture device design and assembly

A custom acrylic frame with inserts for a 6-well tissue culture plate, magnets, and stepper motors was designed to suspend the culture plate above rotating magnets, which in turn were affixed to software-controlled motor shafts throughout culture. The components were designed as a series of two-dimensional panels (Fig. S1, Supplementary Data) in the vector graphics software Inkscape (Inkscape Project), which were laser cut (ILS9.75, Universal Laser Systems) from clear cast acrylic (0.125 in thickness) and assembled with stainless steel socket cap screws. Nema 8 stepper motors (8HS11-0204S and 8HS15-0604S, OMC StepperOnline) with a 1.8° step size were controlled using an Arduino Uno R3 microcontroller board (A000066, Arduino) equipped with an Adafruit Motor/Stepper/Servo Shield v2.3 (1438, Adafruit Industries). All software coding was performed within Arduino IDE v1.8.33.0 (Arduino Software). Cylindrical neodymium magnets (D8A, K&J Magnetics) were fixed to the ends of the motor shafts, providing *ca*. 1 mm of clearance beneath the culture plate with the device fully assembled. All components were sterilized by spraying with 75% ethanol and transferred into a tissue culture incubator set to 37 °C and 5% CO2. One day prior to culture, 6-well (4.67 cm^2^ growth area) Transwell® cell culture inserts (3450, Corning) were coated with 2 mL of 10 µg mL^-1^ human type 1 collagen (5007, Advanced Biomatrix) in 1× PBS (46-013-CM, Corning) for ≥2 h within the same CO2 incubator [39-42]. The inserts were rinsed with 2 mL of 1× PBS immediately prior to cell seeding.

### 2.2. Magnetic agarose hydrogel formation

Deionized (DI) water (≥17.8 MΩ·cm), purified through a Barnstead NANOpure Diamond (Thermo Scientific) or a Milli-Q® IQ 7000 (Millipore-Sigma) filtration system, was used for reagent preparations. Magnetic nanoparticles (MNPs) were synthesized by mixing 23.82 g Fe(II)Cl2•4H2O (AC205082500, Acros Organics) and 38.94 g Fe(III)Cl3 (ICN15349980, MP Biomedicals) in 3 L of DI water. Once dissolved, 240 mL of NH4OH solution (05002, Millipore-Sigma) was added under constant stirring. After 5 min, the formed MNPs were sequentially suspended, magnetically separated, and decanted ≥3 times using 1.5 L of DI water. The final purified MNPs were suspended in DI water at a concentration of >40 mg mL^-1^ and stored within amber bottles at room temperature (20-22 °C) for up to 1 y.

To generate magnetic agarose, a stock solution of 2% w/v agarose was initially generated by mixing the requisite amount of agarose (A9414, Millipore-Sigma) and DI water at 70 °C until dissolved. The liquid stock agarose solution was combined with MNPs and DI water to final concentrations of 1.7% w/v agarose and 10 mg mL^-1^ MNPs. This mixture was stored as a solidified gel at room temperature (20-22 °C) for up to 1 y. To shape the hydrogels for cell culture, the magnetic agarose was melted at 70 °C, stirred, and cast (250 μL of solution) into 8 mm diameter PDMS wells. The magnetic agarose disks were set at room temperature and demolded. The gels were thereafter incubated in 75% ethanol for 5 min and cleansed three times with 50 mL of 1× phosphate-buffered saline (PBS, 46-013-CM, Corning) within a biosafety cabinet under aseptic conditions.

### 2.3. COMSOL modeling of frictional shear forces

COMSOL Multiphysics v5.5a (COMSOL Inc., Burlington, MA) was employed using the Structural Mechanics module to estimate the frictional shear forces applied by a sliding hydrogel. The parameters utilized for the hydrogel were: Young’s modulus (*E*) = 8 kPa (Fig. S3, Supplementary Data), Poisson’s ratio (*ν*) = 0.49 [43], density (*ρ*) = 1010 kg m^3^, and friction coefficient (*µ*) = 0.1 [44]. The parameters utilized for the underlying membrane were: *E* = 2950 MPa [45], *ν* = 0.37 [46], and *ρ* = 1375 kg m^3^ [45]. The simulated gel was placed in conformal contact with the underlying membrane, sliding across the surface at a velocity of 12 mm s^-1^. The average kinetic friction output is relative to the direction of movement and converges to a constant value by *t* = 2500 ms (Fig. S2, Supplementary Data), from which all data are reported in absolute terms.

### 2.4. Shear stress exposure to primary colonic epithelial cells

The human colonic epithelial biopsy specimen (male, 52 y) was obtained during a routine screening colonoscopy performed at the University of North Carolina (UNC) Hospitals Meadowmount Endoscopy Center under UNC IRB #14-2013. The epithelial cells were expanded following a previously reported monolayer culture protocol (RRID:CVCL_ZL23, Supplementary Data) [32, 34]. For seeding into the collagen-coated inserts, cells from one well of a 6-well maintenance/expansion plate (9.6 cm^2^ well^-1^, ≥80% confluency) were split to six collagen-coated Transwells (4.67 cm^2^ insert^-1^). These cultures were expanded to 100% confluency for 4-6 d within an expansion medium (EM) containing Wnt3a, r-spondin 3, and noggin (Table S1, Supplementary Data), before differentiation for 4 d within a differentiation medium (DM) that was absent in these growth factors (Table S1, Supplementary Data). Both EM and DM were replenished every 48 h, with 2 mL of medium added to each luminal and basal compartment. Mechanical stimulation was initiated simultaneously with the switch to DM, with one sterile magnetic agarose disk placed in the luminal compartment of each cell culture insert. The static control inserts were also differentiated under DM for 4 d but did not receive magnetic agarose disks. All companion plates were secured in their respective mechanical culture devices and the appropriate rotation program was initiated through the Arduino microcontrollers until cellular assay or fixation.

### 2.5. In situ assessment of barrier integrity

Transepithelial electrical resistance (TEER) was measured with an EVOM^2^ epithelial volt/ohm meter (World Precision Instruments, Sarasota, FL). Before use, the probe was sterilized in 75% ethanol and rinsed in DM. The culture plates were measured immediately after removal from the CO2 incubator. The resistance was measured in Ohms (Ω) at 3 equidistant points for each culture insert and the values were averaged. A non-seeded, collagen-coated insert remained immersed in medium at 37 °C as a blank control, from which the resistance was also measured and subtracted from the cell-containing inserts each day. Measurements were taken on day 4 of culture, before the cells were differentiated, and on day 8, immediately before LIVE/DEAD staining. Blank-subtracted resistance values were normalized to the insert area (4.67 cm^2^) and the final TEER change between day 4 and 8 values are reported as a percent of the initial day 4 value [47].

### 2.6. Measurement of MUC2 and IL-8 protein secretion

After 4 d of magnetic agarose culture, medium was collected from the luminal and basal reservoirs, and immediately stored at -20 °C for future enzyme-linked immunosorbent assay (ELISA). The medium in the luminal compartment was gently mixed and washed over the cells to ensure all mucins were collected. IL-8 concentrations in medium samples from the basal compartment were measured using the IL-8 Human Uncoated ELISA Kit (88-8086-88, Invitrogen). Assays were performed in Nunc MaxiSorp (tm) flat-bottom 96-well plates (Invitrogen, 44-2404-21) according to the manufacturer’s instructions. The samples were measured using a SpectraMax M5 plate reader (Molecular Devices, San Jose, CA) in fluorescence mode.

Relative amounts of MUC2 in apical medium samples were measured by a separate ELISA. 100 µL of medium (diluted in 0.1 M sodium carbonate, pH 9.5) were loaded into separate wells of a 96-well high affinity ELISA plate (PI15042, Fisher) and dried overnight at 40 °C. The wells were washed with three times with PBS and blocked with 1% BSA (A9647, Sigma) for 1 h. Horseradish peroxidase (HRP)-conjugated MUC2 detection antibody (sc-515032, Santa Cruz Biotechnology) was diluted to 15 ng mL^-1^ in 1% BSA, and 100 μL of this solution was added to each well. The plates were incubated in this solution at 4 °C overnight. The wells were washed five times with 0.05% Tween-20 in PBS, and 100 µL of SuperSignal ELISA Femto Substrate detection solution (Fisher, PI37075) was added to each well and incubated at room temperature (20-22 °C) for 20 min. The samples were measured using a SpectraMax M5 plate reader (Molecular Devices, San Jose, CA) in luminescence mode.

### 2.7. Fluorescence microscopy

The cells were assayed by fluorescence imaging of DNA (Hoechst 33342, B2261, Millipore-Sigma, and propidium iodide, P3566, Life Technologies), intracellular esterase activity (354217, Corning), alkaline phosphatase (ALP) activity (SK-5100, Vector Laboratories), mucin-2 (MUC2) presence (sc-515032 AF488, Santa Cruz Biotechnology), actin localization (R37110, ThermoFisher Scientific), E-cadherin localization (sc-7870, Santa Cruz Biotechnology), and zonula occludens-1 (ZO-1) localization (21773-1-AP, Proteintech Group) using established labeling methods (Supplementary Data) [34, 41, 42]. Confocal fluorescence microscopy was performed on an inverted Olympus Fluoview 3000 (Waltham, MA) equipped with 405, 488, 561, and 640 nm laser diodes operating in conjunction with a galvanometer scanner. Emission wavelengths (Hoechst: 430-470 nm, Cy5: 650-750 nm, AF 488: 505-545 nm, Texas Red: 600-640 nm) were selected from manufacturer-provided control software. For the LIVE/DEAD, MUC2, and ALP assays, the entire Transwell insert was mounted over a glass coverslip in 1× PBS and imaged as an 8×8 tiled image grid (3.11 µm px^-1^, 1024 px^2^ area, 15% overlap) using a 4× objective (N.A. 0.16, UPlanSApo4X). The images from each Transwell insert were automatically stitched during acquisition using the manufacturer-provided control software. For each region of interest (ROI), at least 3 *z*-series optical sections were collected with a step size of 25 µm to account for sample thickness. For the remaining target proteins (F-actin, E-cadherin, ZO-1), the samples were mounted over a glass coverslip in a fructose/glycerol clearing solution [48] and imaged as separate ROIs using a 30× silicone oil immersion objective (N.A. 1.05, correction collar: 17 mm/23 °C, UPlanSApo30XS) and 2× digital zoom (0.21 µm px^-1^, 1024 px^2^ area). To account for potential regional variations in cellular morphology, 6 images per Transwell insert were acquired at equidistant points along the magnetic hydrogel path. For each ROI, at least 40 *z*-series optical sections were collected with a step size of 0.75 µm.

### 2.8. Image analysis

Image areas positive for LIVE/DEAD markers, mucin-2, and alkaline phosphatase reaction precipitate were quantitatively evaluated and normalized to nuclear area or total insert area, as specified below. *Z*-series optical stacks were flattened into two-dimensional representations through maximum intensity *z*-projections (Calcein-AM and propidium iodide) or median intensity z-projections (MUC2 and ALP), with brightness/contrast and window/level adjusted linearly and identically for compared image sets in cellSens Dimension v1.18 (Olympus Corp.) and the Fiji software package [49]. Within Fiji, each fluorescence channel was converted to binary through empirical threshold selection. The average area occupied by the suprathreshold level of fluorescence was measured and exported to a spreadsheet. Extracted suprathreshold area values were normalized to Hoechst 33342^+^ area (for MUC2 and ALP), total insert area (for calcein-AM) or calcein-AM^+^ area (for propidium iodide).

Morphological feature extraction utilized ZO-1 demarcated cell boundaries and a semi-automated image analysis pipeline, while nuclear feature extraction utilized Hoechst 33342^+^ areas and a fully automated image analysis pipeline (Supplementary Data). For the ZO-1 boundary images, *Z*-series optical stacks were subjected to three-dimensional constrained iterative deconvolution in cellSens Dimension v1.18 (Olympus Corp.) utilizing 5 iterations of an advanced maximum likelihood algorithm, with automatic denoising, background removal, and edge protection options enabled. The deconvolved images were flattened into two-dimensional representations through maximum intensity *z*-projections, and thereafter imported into CellProfiler v3.1.9 for further processing and segmentation [50]. Each two-dimensional image was flat-field corrected using a gaussian-smoothed illumination correction function (150 px filter size). The corrected images were converted to binary using 3-class Otsu threshold selection, with an adaptive window of 20 px and the middle pixel intensity class assigned to the background. The thresholded images were thinned to single pixel wide morphological skeletons, which were infinitely cleaned to remove isolated pixels before a morphological closing operation was applied using a disk structuring element (5 px diameter) to connect any incomplete paths. The final skeleton images were inverted, eroded with a disk structuring element (1 px diameter), and holes (<5 px diameter) were filled to remove any remaining noise. A distance-dependent watershed segmentation algorithm with a footprint of 60 px was applied for object identification. All objects were property filtered by location (border objects removed) and Euler’s number (*e* < 1 removed), with a final manual intervention step incorporated for quality control of over-segmented (7 ± 6 cells image^-1^, requiring join operation), under-segmented (4 ± 6 cells image^-1^, requiring split operation), or misidentified (2 ± 2 cells image^-1^, requiring deletion) cells (Table S2, Supplementary Data). The area, perimeter, circularity (“FormFactor,” 4π × area × perimeter^-2^), solidity, and major/minor axis lengths for the remaining objects were measured and exported to a spreadsheet.

### 2.9. Statistical analysis

Statistical analyses and data plotting were performed in GraphPad Prism v9.0.1 (GraphPad Software, San Diego, CA) at a significance level (*α*) of 0.05 unless otherwise noted. Group means from TEER (*n* = 4 Transwells condition^-1^) and LIVE/DEAD staining (*n* = 4 Transwells condition^-1^) measurements were analyzed by 2-way ANOVA followed by Bonferroni post-tests for multiple comparisons. Group means from MUC2 ELISA (*n* = 11 Transwells condition^-1^), IL-8 ELISA (*n* = 4 Transwells condition^-1^), MUC2 immunofluorescence (*n* = 5 Transwells condition^-1^), and ALP immunofluorescence (*n* = 5 Transwells condition^-1^) measurements were analyzed by unpaired, two-tailed t-tests. Welch’s correction was applied for the MUC2 ELISA comparison to account for unequal variances between datasets (F-test, *p* < 0.001).

Cell morphology data analysis was performed in RStudio v1.3.1056 (RStudio Team, Boston, MA) running R x64 v4.0.2 (R Core Team, Vienna, Austria) [51, 52]. SuperPlots were generated by pooling the means of the morphological parameters from each replicate Transwell insert, reducing the sample size from the total number of cells (n > 5000 cells condition^-1^) to the number of technical replicates (*n* = 5 Transwells condition^-1^), followed by an unpaired, two-tailed t-test to determine *p* values [53].

Unless specified, data are presented as sample means with error bars depicting one standard deviation. The box plot crossbars represent sample medians, 25^th^ percentiles, and 75^th^ percentiles, with the whiskers depicting the minimum and maximum measurements from each dataset. For statistical comparisons, *p* values are represented as follows: * for *p* ≤ 0.05, ** for *p* < 0.01, and *** for *p* < 0.001.

## 3. Results and discussion

### 3.1. Generation of a solid-induced shear force model for primary colonic epithelial cells

Agarose was selected as the analog for solid fecal matter as it is widely available, exhibits facile molding/handling capabilities, possesses a controllable stiffness that can be tailored within the known range of fecal material, and is known to be compatible with primary tissue culture [54]. Among several fabrication conditions tested, an agarose concentration of 1.7 mg mL^-1^ demonstrated the most consistent molding capabilities, providing a freestanding structure that could be manipulated using forceps and subjected to solvent sterilization procedures. Propulsion of the hydrogel over the epithelial surface was noninvasively actuated *via* a neodymium magnet secured beneath the culture plate. To render the hydrogels magnetically active, superparamagnetic iron oxide nanoparticles were embedded within the agarose at a concentration of 10 mg mL^-1^ (Fig. 1a). These magnetic hydrogels were movable in a direction parallel to the epithelial surface when the neodymium magnet was moved parallel to the plate surface. Hydrogels with higher nanoparticle concentrations remained parked in a single location when the magnet was rotated below the plate due to an excessive downward force creating an insurmountable frictional force, while those with lower nanoparticle concentrations could not be reliably rotated across the epithelium due to an insufficient forward magnetic force. The measured storage modulus of magnetic hydrogel was 2.6 ± 1.2 kPa (*n* = 5 hydrogels, Young’s modulus *ca*. 7.8 kPa, Fig. S3). A computational model, consisting of a hydrogel of this stiffness placed in conformal contact with an underlying membrane, was created to better understand the frictional shear forces imparted to the underlying epithelial cell surface during lateral propulsion. This model operated under the assumption that the contribution of downward magnetic pull force is negligible compared to the applied load from gravity. When the simulated gel slides across the underlying surface at a rate of 12 mm s^-1^, the model predicted an average kinetic friction of 4.95 Pa (*t* = 2500 ms, Fig. 1c and S2). These results follow Amontons’ first law in that the frictional force is directly proportional to the applied load. Therefore, the shear forces can easily be modulated by users of this platform through simple tuning of the hydrogel size/weight, depending on the experimental design and hypotheses being tested.

**Fig. 1.**
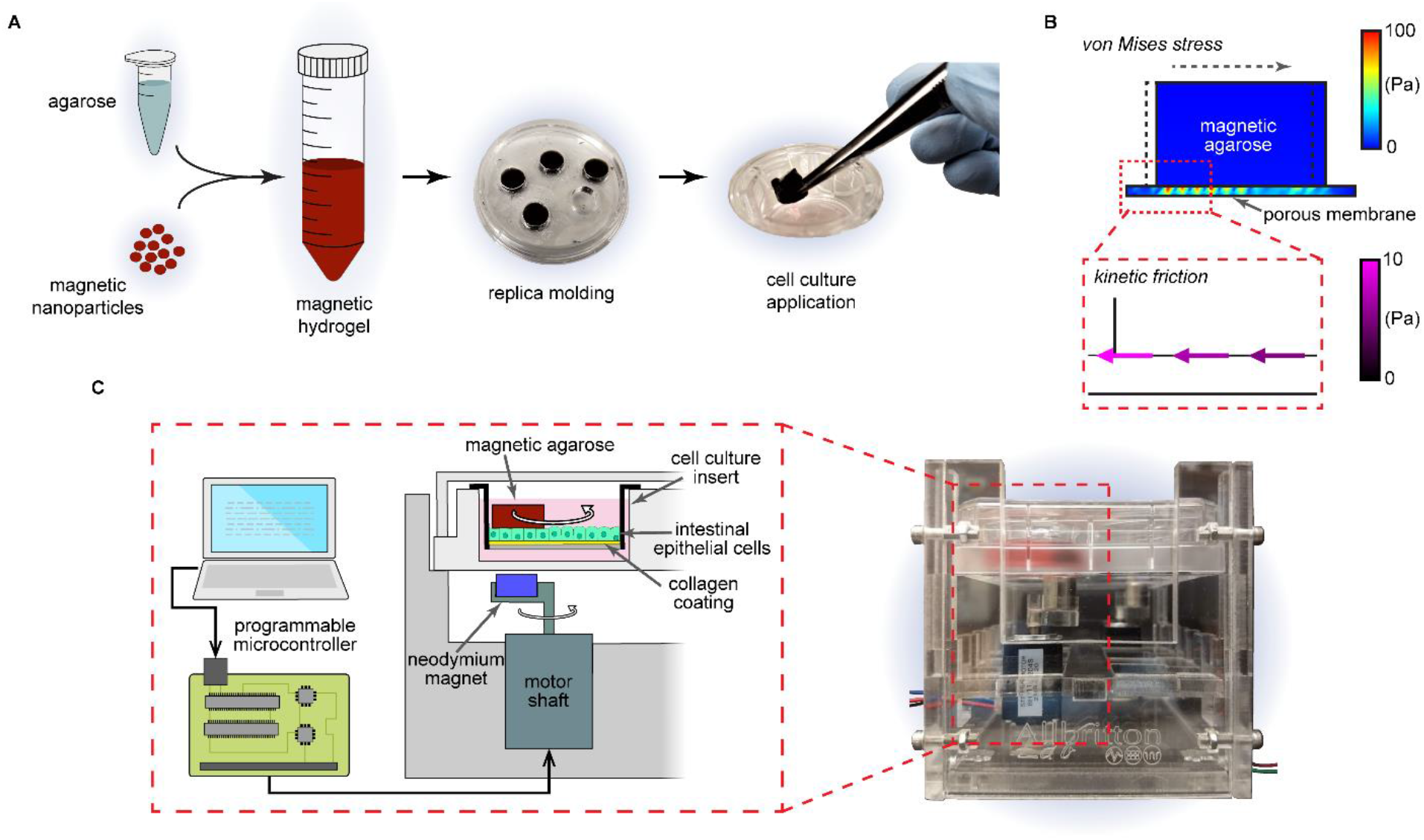
Magnetic agarose fabrication and device assembly. **(A)** The magnetic agarose is formed by mixing of liquid agarose and suspended magnetic nanoparticles. This mixture is set into a stable gel at ambient temperature, solvent sterilized, and applied to primary human intestinal epithelial cells. **(B)** COMSOL modeling of von Mises stress and kinetic friction as the magnetic agarose is propelled counterclockwise across an underlying membrane surface (from left to right in the stress plots). The magnetic agarose experiences kinetic friction in the opposite direction of its movement. The direction and magnitude of this force at every 1.33 mm are indicated by the purple gradient arrows in the kinetic friction inset. The results shown are from *t* = 2500 ms, after convergence of average values. **(C)** Schematic of the programmable microcontroller, magnetic motor shaft, and cell culture interfaces, with photograph of the fully assembled device.

Commercial microcontrollers and standard cell culture plates were incorporated into the device design to maintain compatibility with established assay infrastructure, facilitate automation, and enable parallel experimentation. The neodymium magnets beneath the plate were affixed to the shafts of computer-controlled motors, enabling programmable alterations of the frequency, duration, and speed of the magnetic hydrogel movement. Microstepping of the motors (1.8° step size) provided smooth motion during periods of rotation instead of stepwise movements (Video 1, Supplementary Data), and a custom frame was constructed from laser-cut acrylic sheets to interface all device components throughout culture (Fig. 1d and S1, Supplementary Data). The specific materials and design ensure standardized positioning and manipulation of the neodymium magnets relative to the magnetic agarose and can be readily adapted for various plate sizes and manufacturing methods.

### 3.2. Optimization of device parameters for primary cell compatibility

Primary human colonic epithelial stem cells were expanded from biopsied tissue by isolating the colonic crypts and culturing the cells as a monolayer within a growth factor-enriched medium [30] over a soft collagen hydrogel scaffold [32, 34]. Compared to the widely used organoid culture technique, this hydrogel culture method permits rapid scale-up and facile serial passaging of intestinal epithelial cells. Additionally, loss of genetic integrity has not been observed in the monolayer cultures for up to 15 passages [34]. These neutralized collagen scaffolds proved to be incompatible with magnetic agarose hydrogel propulsion over their surface since their low stiffness resulted in permanent deformation or damage by the applied shear forces (Fig. S4, Supplementary Data). To promote long-term stability of the platform under repeated shear stress over time, a porous membrane with a surface coating of extracellular matrix (ECM) was adopted for cellular support. We have previously demonstrated hanging basket insert systems with basal porous membranes supply critical nutrients and growth factors to the basal tissue side and support a healthy and functional cell monolayer that is representative of primary intestinal tissue (*e.g*., proliferative regions, differentiated lineages, transport, and barrier function) [35-37, 42]. The epithelial monolayer also establishes proper intercellular focal adhesions creating a high resistance barrier that segregates the luminal and basal culture compartments, enabling measurements of directional transport or secretion. The system is well-suited to *in situ* assays of living cells over time since the monolayer is easily accessed and also supports endpoint assays of fixed cells. Importantly, given the stiffness of the ECM-coated membranes, these surfaces with attached cells were likely to resist mechanical deformation by magnetic agarose propulsion over the surface (Video 1, Supplementary Data).

To generate the shear force exposure model, colonic epithelial cells were first seeded over ECM-coated porous membranes and cultured under an expansion medium (containing WNT3a, r-spondin 3, noggin, and epidermal growth factor) until the intestinal monolayers became fully confluent over the luminal membrane surface (4 – 6 d). Once contiguous monolayers of proliferative cells were formed, differentiation into absorptive and secretory lineages, which comprise >90% of colonic epithelial cells *in vivo* [3], was induced by removal of the above growth factors (*i.e*., culture in differentiation medium) and shear forces were introduced by application of the magnetic agarose hydrogels. Propulsion of the hydrogels over the epithelial surface was actuated at a set frequency and linear velocity for 4 d, at which time LIVE/DEAD staining assays were utilized to determine the range of parameters that were compatible with colonic epithelial viability (Fig 2). The device parameters tested included linear velocities of 0 – 40 mm s^-1^ and rotation frequencies of 0 – 12 events h^-1^. Increasing the linear velocity of the fecal surrogate from 0 to 40 mm s^-1^ did not affect cellular viability or surface coverage (Fig 2b,c). While no differences were noted between the control cultures and those exposed to shear forces at a rotation frequency of 1 event h^-1^ (60 min interval between rotations), increasing the frequency to 12 events h^-1^ (5 min interval between rotations) resulted in a lower cellular surface coverage and increased cell death at 2 mm s^-1^. This may be the result of a much higher contact time between the magnetic agarose and the luminal face of epithelial cells compared to the other conditions tested, as well as increased static friction during the initiation of magnetic agarose propulsion. Qualitatively, epithelial monolayers cultured with a magnetic agarose rotation frequency of 12 events h^-1^ displayed vacancies of cells in which large regions of the monolayer were detached. However, this was not statistically reflected in the live cell area quantitations due to a much higher measurement variability under the high frequency conditions (Fig 2b).

**Fig. 2.**
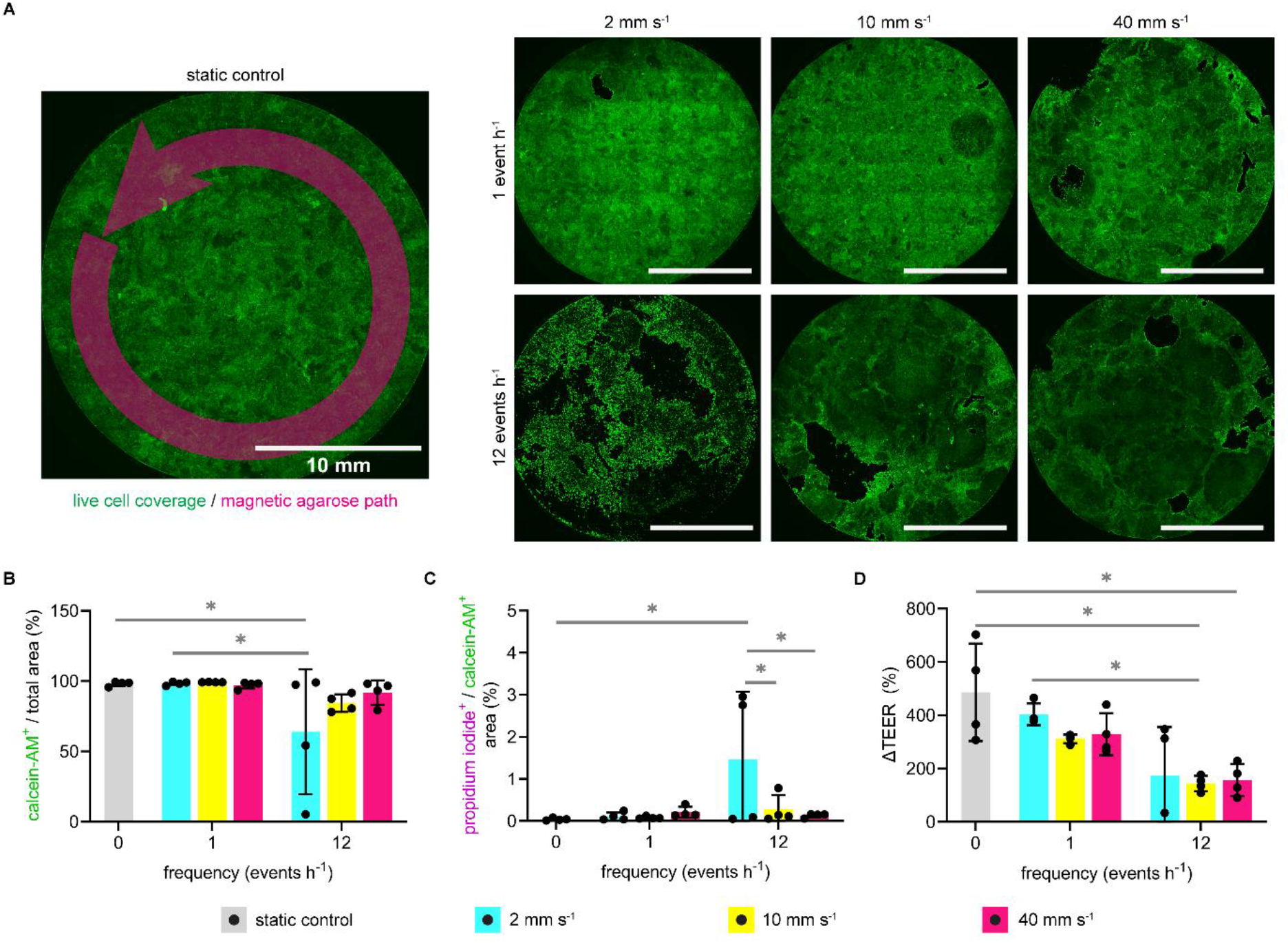
Characterization of device parameters operating with primary human colonic epithelial cells. **(A)** Fluorescence microscopy images of calcein-AM treated cells (green) under magnetic agarose propulsion with varied linear velocities (0 – 40 mm s^-1^) and rotation frequencies (0 – 12 events h^-1^). The path of the magnetic agarose is highlighted in magenta. Scale bars represent 10 mm. **(B)** Effects on live cell coverage, assayed by calcein-AM fluorescent staining (*n* = 4, 2-way ANOVA, Bonferroni post-test). **(C)** Effects on cellular viability, assayed by propidium iodide fluorescent staining (*n* = 4, 2-way ANOVA, Bonferroni post-test). **(D)** Effects on transepithelial electrical resistance (*n* = 4, 2-way ANOVA, Bonferroni post-test).

Transepithelial electrical resistance (TEER) measurements were utilized to assess barrier integrity of the monolayers during optimization of the device parameters (Fig. 2d). The resistance across the monolayers was measured before and after 4 d of magnetic agarose rotations under differentiation medium. It is expected that as the intestinal stem/proliferative cells differentiate to form a mature epithelium, TEER will increase with a larger positive change over the monitoring period being indicative of an intact and mature monolayer possessing high barrier integrity. Analagous to the surface coverage and viability assays, no significant differences in TEER were noted between the control cultures and those exposed to shear forces at a frequency of 1 event h^-1^. However, increasing the rotation frequency to 12 events h^-1^ at 2 – 40 mm s^-1^ resulted in lower TEER measurements compared to that of the control cultures, suggesting that shear force frequencies of this magnitude have a detrimental effect on intestinal barrier function. Taking the TEER and viability data together with established human physiological benchmarks for high amplitude peristaltic waves, a linear velocity of 10 mm s^-1^ and rotation frequency of 1 event h^-1^ were selected for all subsequent investigations.

### 3.3. Differentiated lineage allocation is not affected by frictional shear forces

Lineage allocation of differentiated intestinal epithelial cells into absorptive and secretory phenotypes drives overall function and impacts many pathological responses of the *in vivo* tissue [3]. Previous work using drosophila models suggests that mechanically sensitive ion channels are involved in modulating intestinal stem cell proliferation and directing the differentiation process [9]. While the behavior of cultured mammalian intestinal stem cells is sensitive to mechanical variations of the extracellular matrix [2, 6], the application of periodic frictional shear stress and its downstream effects on differentiation have not been evaluated using primary human intestinal epithelium. After 4 d of magnetic agarose rotations at 10 mm s^-1^ and a frequency of 1 event h^-1^, epithelial cells were fixed and assayed for alkalkine phosphatase (ALP) activity and intracellular mucin-2 (MUC2) presence, which are common biomarkers for absorptive enterocytes [55, 56] and mucus-producing goblet cells [57], respectively (Fig. 3a). Quantification of these fluorescent markers demonstrate that neither is significantly modulated (Fig. 3b), suggesting that the application of shear forces at a velocity and frequency that is representative of healthy human tissue does not affect stem cell fate decisions with regard to goblet or enterocyte lineage allocation. These results are consistent with those obtained from Caco-2BBE cells exposed to shear stress [58], yet are in contrast with those obtained from Caco-2 and HT29-MTX cells exposed to varying mechanical stresses [28, 59, 60]. Given the status of the above lines as heterogenous tumor cell populations with vastly different functional properties than healthy colonic tissue [33, 61-63], models derived from primary healthy colonic epithelial stem cells may be more predictive for understanding the impact of shear forces *in vivo*.

**Fig. 3.**
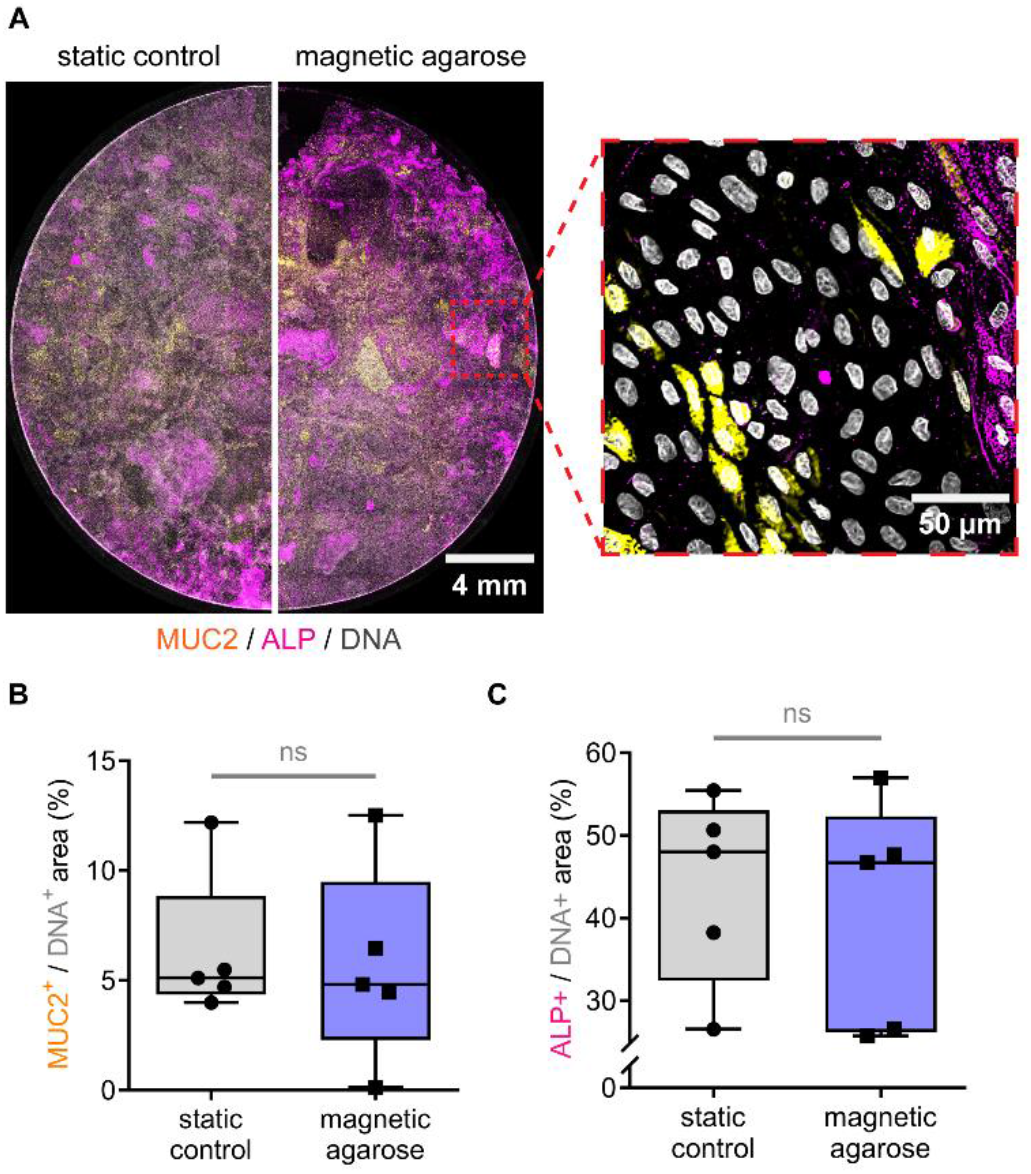
Assay of mucin-2 (MUC2) presence and alkaline phosphatase (ALP) activity by fluorescence microscopy. **(A)** Fluorescence microscopy images of colonic epithelial cells assayed for MUC2 presence (yellow), ALP activity (magenta), and DNA presence (grey) as markers for goblet cells, absorptive colonocytes, and nuclei, respectively. **(B)** Comparison of MUC2^+^ area normalized to DNA^+^ area between static control cultures and magnetic agarose exposed cultures (*n* = 5, *p* = 0.8097, t-test). **(C)** Comparison of ALP^+^ area normalized to DNA^+^ area between static control cultures and magnetic agarose exposed cultures (*n* = 5, *p* = 0.7194, t-test).

### 3.4. Modulation of mucin-2 and interleukin-8 secretion

Shear stresses and luminal pressure variations within the gastrointestinal tract have been linked to the modulation of signal transduction pathways that affect protein secretion by the epithelium [64]. Though the differentiation of intestinal stem/proliferative cells into mucus-producing goblet cells was unaffected by magnetic agarose propulsion (Fig. 3b), the secretion of mucus by the existing goblet cell population might be influenced by mechanical stimulation as a homeostatic or protective mechanism [28, 65, 66]. Therefore, mucin-2 (MUC2) that was secreted into the luminal compartment over the final 48 h of magnetic agarose culture was sampled and measured by ELISA (Fig. 4a). Consistent with studies in which cultured cells were exposed to shear forces [28, 65], MUC2 production increased by nearly two-fold when the epithelial surface was exposed to magnetic agarose rotations relative to that of static control cultures. Increased MUC2 secretion in response to mechanical stimulation may be useful *in vivo* for increasing lubrication and decreasing damage from the propulsion of solid fecal contents [65]. This is supported by measurements of mucus accumulation rates in rat models, which were highest in the colon compared to all other segments of the gastrointestinal tract [66].

**Fig. 4.**
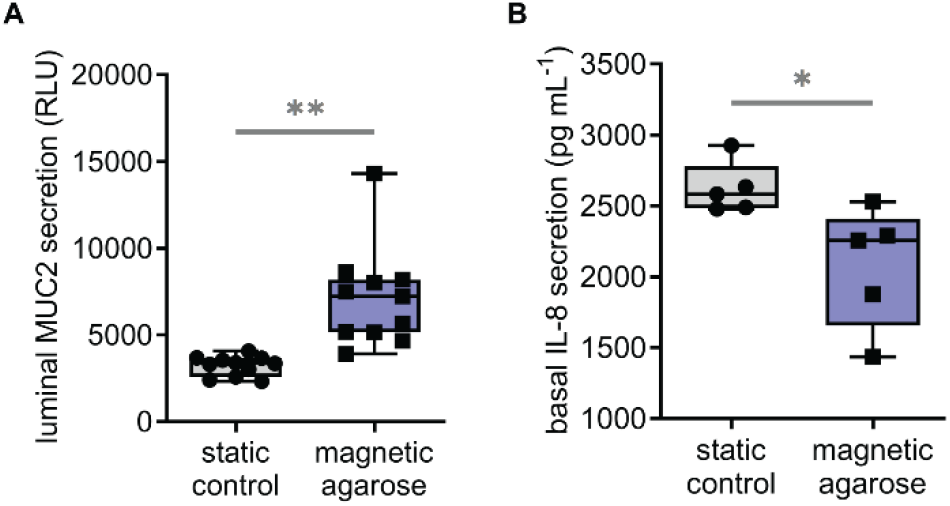
Protein secretion modulation by luminal shear forces. **(A)** Comparison of mucin-2 (MUC2) secretion between static control cultures and magnetic agarose exposed cultures (*n* = 11, *p* = 0.001, Welch’s t-test). (B) Comparison of interleukin-8 (IL-8) secretion between static control cultures and magnetic agarose exposed cultures (*n* = 5, *p* = 0.0307, t-test).

Interleukin-8 (IL-8) is an inflammatory cytokine produced by the intestinal epithelium as part of the response system to pro-inflammatory molecules or bacterial invasion [35, 67, 68]. Acting as a chemoattractant for phagocytic immune cells deep within the *lamina propria* [69-71], induction of directional IL-8 secretion has been observed in several models of primary intestinal epithelium in response to pro-inflammatory insult [35, 72]. While the luminal secretion of IL-8 in these culture systems is consistently high (>5000 pg mL^-1^) and insensitive to inflammatory mediators, the basal concentration is relatively low and readily modulated [35]. Culture medium was therefore collected from the basal reservoir of the platform over the final 48 h of culture and IL-8 was measured by ELISA (Fig. 4b). Intriguingly, basal IL-8 secretion decreased in response to the luminal shear stresses imparted by magnetic agarose rotation. Beyond serving as a general indicator of health for cells cultured within this platform, mechanosensitive modulation of cytokine secretion has been suggested by other reports though not directly assayed [10, 25]. The presence of an immune surveillance mechanism that is modulated by the presence and movement of feces and bacteria may explain this behavior and warrants further investigation of underlying signaling pathways.

### 3.5. Several cellular morphological parameters are impacted by solid-induced shear forces

The epithelial monolayers were subjected to morphological analysis of several key structural proteins to assess their proper localization and basolateral polarity after exposure to shear stress. Actin is concentrated within the microvilli of mature intestinal epithelial cells *in vivo*. This feature was recapitulated for both the static control and shear force exposed cells, which exhibited actin-enriched apical brush border membranes (Fig. 5a). Compared to the static control cultures that exhibit flat luminal surfaces, a more rounded luminal surface can be observed for the cells exposed to shear forces, though average cell heights remain unchanged at 19.0 ± 4.9 µm (*n* = 47 cells, 3 technical replicates) for the static control cells and 16.4 ± 3.9 µm (*n* = 28 cells, 3 technical replicates) for the shear force-exposed cells (*p* = 0.58, Fig. S5). Representative tight junction proteins, zonula occludens-1 (ZO-1) and E-cadherin, are also present in both conditions and with their proper boundary localizations (Fig. 5a). Overall, the intestinal epithelial monolayers exposed to magnetic agarose did not exhibit cytoskeletal disruption. Moreover, when these results are taken together with the TEER measurements, this platform is suitable for transport studies that require epithelial monolayers of high barrier integrity, as is the case for investigations of food, bacterial metabolite, or drug transport as well as directionally secreted proteins.

**Fig. 5.**
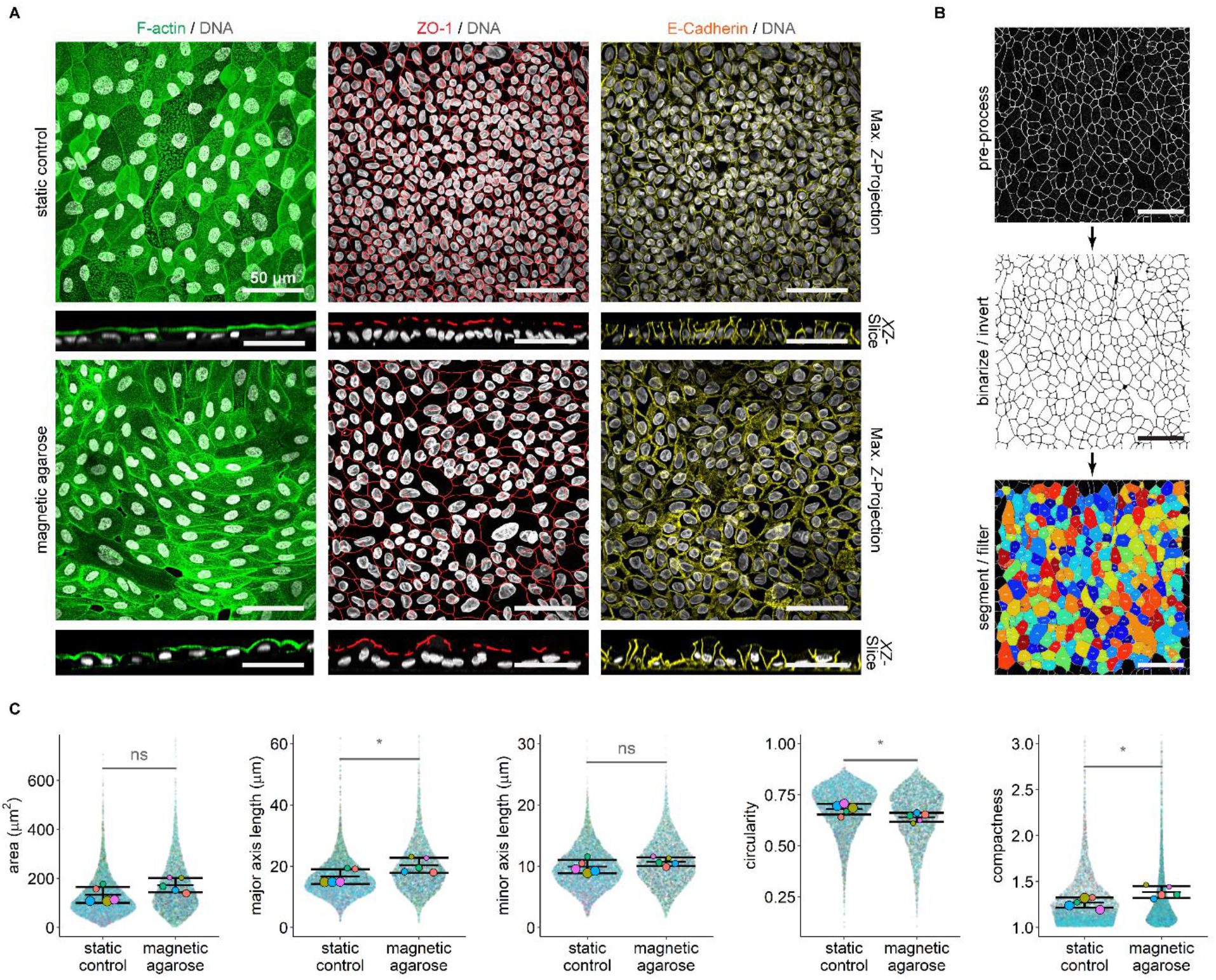
Impacts on cellular morphology. **(A)** Fluorescence microscopy images of colonic epithelial cells assayed for the following markers: F-actin (green), zonula occludens-1 (ZO-1, red), E-cadherin (yellow), and DNA (grey). Scale bars represent 50 µm. **(B)** Summary of image processing pipeline for extraction of morphological features from ZO-1 labeled fluorescence microscopy images. Scale bars represent 50 µm. **(C)** Comparisons of cellular area (*p* = 0.074), major axis length (*p* = 0.043), minor axis length (*p* = 0.2), circularity (*p* = 0.031), and compactness (*p* = 0.017) between static control cultures and magnetic agarose exposed cultures (*n* = 5, t-test). The transparent data points represent individual cells (*n*_static control_ = 7927 cells, *n*_magnetic agarose_ = 5789 cells) and are color-coded by technical replicate, while the solid data points represent the mean value for each technical replicate and are sized by the number cells identified in each hanging basket insert (*n* = 5). The mean crossbar, error bars, and t-tests were calculated from the solid data points.

While the epithelial monolayers remained fully intact under magnetic agarose propulsion, with no fragmentation or diminished expression of structural proteins, the tight junctions of these cells displayed alterations in cellular morphology, appearing more elongated and irregular in shape, and often in alignment with the magnetic agarose path, compared to the control cells (Fig. 5a). To parse the more subtle effects of shear force exposure on lateral cellular morphology, changes to the tight junction protein, ZO-1, including its location and patterning, were quantitatively assessed using a customized image analysis pipeline (Fig. 5b). As ZO-1 is restricted to the luminal face of the epithelial cells and is thus closest to the source of the applied shear forces, this tight junction protein provides excellent representation of the morphological response of the cells. While each of the morphological parameters exhibited measurable mean differences upon shear force exposure (Fig. 5c), only the circularity, compactness, and major axis lengths demonstrated significant changes (at a threshold of *α* = 0.05) when the cells were aggregated by technical replicate (Fig. 5c and Table S3, Supplementary Data). The decrease in circularity and increase in major axis length (but not minor axis length) are consistent with the cells appearing more elongated and are also present in observed nuclear morphology trends (Table S4, Supplementary Data). Histologic analysis of the luminal surface cells from human colorectal tissue have demonstrated higher nuclear axis ratios (major : minor axis lengths) and lower nuclear circularity indices compared to basal crypt cells [73]. The presence of luminal shear forces may assist in the development of this surface epithelial cell morphology *in vivo*, as suggested by our nuclear and ZO-1 cell boundary analysis. Meanwhile, the observed increase in compactness indicates that the cells are less regular in shape with jagged or indented borders (Fig. 5c). This may be an early sign of luminal exfoliation, which has previously been characterized by marked irregularities in the plasma membrane [74]. There is no transition of the cells to a flattened or squamous state that can be representative of a cancerous phenotype [75], as the area, perimeter and height are minimally affected.

## 4. Conclusions

Mechanical stimulation by solid fecal contents is a widely overlooked area of colonic epithelial physiology. All previously developed intestinal culture platforms have sought to characterize mechanical stimulation through stretch-strain of the underlying basal support or through luminal liquid microflow, with most of these systems using tumor cell lines that are not fully representative of *in vivo* human tissue. While studies utilizing primary cells grown in some of these microfluidic platforms may shed light on how low-amplitude propagated contractions of the colon affect epithelial homeostasis [27], the effects of high-amplitude propagated contractions and the luminal propulsion of solid fecal contents have not been recapitulated. To investigate this aspect of human physiology, we have designed a platform that mimics fecal propulsion over the colonic epithelium through noninvasive actuation of a magnetic agarose hydrogel. The device components, including software-controlled magnetic stepper motors and a customized culture plate housing, were designed around commercially available cell culture insert systems possessing standardized spacing to facilitate automation and compatibility with existing assay infrastructure. Such platforms can be readily adopted by standardly equipped biomedical research laboratories using easily procurable materials and assembly methods for rapid integration into established experimental pipelines. We also demonstrated the ease of sampling the basal/luminal culture medium for analytical assay and performing *in situ* imaging of both live and fixed cells. As the platform is optimized for primary human colonic epithelial cells, the observed cellular behaviors and responses are likely to be more predictive of human *in vivo* tissue than those observed with tumor cell lines [11, 38].

The magnetic agarose hydrogel that was developed for this study proved sufficiently robust for solvent sterilization, manual manipulation, and magnetic propulsion, yet was compatible with primary cell culture and capable of eliciting cellular responses through shear force induction. Interestingly, the normal physiological frequency and velocity values for high-amplitude propagated contractions for the human colon were within the experimental parameters that exhibited high cell viability and TEER in our system. However, at the highest rotational frequencies and velocities tested, representing a 50× greater contraction frequency and 4× greater fecal velocity than observed in healthy humans [7, 12, 14], large areas of cell monolayer were detached from the culture insert with accompanying decreases in viable cell coverage and transepithelial electrical resistance (TEER). Such results highlight the importance of surveying environmental factors that are present during pathological conditions in which colonic motility is affected, as is the case for clinical patients experiencing slow-transit constipation or chronic diarrhea [14, 20]. Beyond enabling the opportunity to experimentally tune these parameters, future research directions include altering the magnetic hydrogel to match the rheological properties and water contents that are present across the entirety of the Bristol Stool Form Scale [19], which will be supported by the increased availability of rheological data on human feces [15-18].

At a frequency and velocity that match high-amplitude propagated contractions of healthy humans, propulsion of magnetic agarose over the epithelial surface did not affect lineage allocation into absorptive or mucus-producing cell phenotypes, though shear stress modulated both mucin-2 (MUC2) and interleukin-8 (IL-8) secretion. The mechanistic connection between shear force exposure and the secretion of these two proteins may be explained by recent investigations of intestinal Piezo2 mediated serotonin release [10, 25]. Colonic enterochromaffin cells secrete serotonin when mechanically stimulated which is thought to be due to opening of the stretch-activated Piezo2 ion channel [10, 25, 26, 76]. Thereafter, serotonin may locally modulate mucus production by goblet cells [77] or regulate inflammation [78, 79] while promoting peristalsis, vasodilation, and pain sensation [76]. These results agree with our observations of secreted protein modulation after shear force induction. Further investigations that utilize newly developed primary culture techniques in which goblet cell lineages [36] or enterochromaffin cell lineages [37, 80] are enriched will be important platforms when used in combination with the magnetic agarose fecal mimic to understand the role of Piezo2 mediated mechanotransduction in response to shear forces.

Morphological analysis of the epithelial cells exposed to magnetic agarose-induce shear force revealed that critical structural and tight junction proteins were present and in their proper basolateral locations. Compared to previous microfluidic studies [22, 23, 27, 28], we observed no change in cellular height or the formation of villus structures when solid-induced shear forces were applied, although significant alterations were noted in lateral morphology which have not been previously documented. Increases in cellular height leading to highly columnar cells within self-renewing monolayers have recently been ascribed to enhanced oxygen availability [81] which is likely facilitated in microfluidic systems through continuous medium flow. The effects of shear forces on stem/proliferative populations and under varying oxygen tensions merit investigation to understand their holistic contributions to morphology and long-term injury repair cycles [81]. In summary, the platform developed in this report represents an ideal method for examining the broad scope of effects that accompany direct mechanical stimulation as it utilizes primary human cells, enables programmable alterations of colonic motility, and can be readily adapted by others. The results of any future mechanotransduction studies will undoubtedly be bolstered by the inclusion of other critical components of intestinal tissue, including the microbiome, immune cells, and fibroblasts, which will present their own mechanosensitive reactions [82, 83].

## Supporting information

Video 1. Magnetic Agarose Rotation

## ACKNOWLEDGMENTS

We acknowledge financial support from the National Institute of Diabetes and Digestive and Kidney Diseases (NIDDK) at the National Institutes of Health (grant numbers R01 DK109559 and R01 DK120606 to N.L.A.), the National Institute of General Medical Sciences at the National Institutes of Health (grant number R35 GM138036 to C.A.D.), the National Science Foundation (graduate research fellowship 2018261576 to R.C.B.), and the University of Washington (start-up funding). Laser cutting was performed within the Be A Maker (BeAM) network of makerspaces at the University of North Carolina at Chapel Hill. The authors wish to thank Prof. Scott T. Magness for procurement of the intestinal biopsy specimen and Angelo Massaro for device assembly assistance.

## DATA AVAILABILITY

The imaging data and code required to reproduce these results are available to download from: https://dx.doi.org/10.6084/m9.figshare.14036855.

## APPENDIX A. SUPPLEMENTARY DATA

Supplementary data to this article can be found online at […], and includes: Supplementary data file (.docx) and Video 1 (.mp4).

