## Supplementary Data file for "Magnetically-Propelled Fecal Surrogates for Modeling the Impact of Solid-Induced Shear Forces on Primary Colonic Epithelial Cells"

**Materials and Methods** . . . . . . . . S-2

**Tables** . . . . . . . . . . S-5

Table S1. Medium formulations . . . . . . . . S-5

Table S2. Quality control for cell morphology pipeline . . . . . S-6

Table S3. Cellular morphology summary statistics . . . . . S-6

Table S4. Nuclear morphology summary statistics . . . . . S-7

**Figures** . . . . . . . . . . S-8

Figure S1. Acrylic frame schematics . . . . . . . S-8

Figure S2. Time vs. average kinetic friction model . . . . . S-8

Figure S3. Magnetic agarose rheology . . . . . . . S-9

Figure S4. Deformation of neutralized collagen by magnetic agarose . . . S-10

Figure S5. Comparison of cellular heights . . . . . . S-11

**References** . . . . . . . . . . S-12

### Materials and Methods

*Buffer Compositions*

DI water (≥17.8 MΩ·cm), purified through a Barnstead NANOpure Diamond (Thermo Scientific) or a Milli-Q® IQ 7000 (Millipore-Sigma) filtration system, was used for all reagent preparations. *Phosphate buffered saline (1× PBS)*: 10% (v/v) Corning® 10× PBS (46-013-CM, Corning), pH 7.4. *Trizma buffer:* 7.92 mg $mL^{-1}$ Trizma HCl, 12.1 mg $mL^{-1}$ M Trizma Base, pH 8.4. *Immunofluorescence (IF) buffer:* 10% (v/v) 10× PBS, 0.2% (v/v) Triton X-100, 0.05% (v/v) Tween-20, 1 mg $mL^{-1}$ bovine serum albumin, 0.5 mg $mL^{-1}$ sodium azide.

*Monolayer Culture and Expansion of Primary Human Colon Epithelial Cells*

Whole intestinal crypts were isolated and expanded as outlined previously [1, 2]. Briefly, crypts were obtained from a biopsy (male, 52 y) collected during a routine screening colonoscopy performed at University of North Carolina (UNC) Hospitals Meadowmount Endoscopy Center with consent of the patient under UNC IRB #14-2013 (RRID:CVCL_ZL23). Crypts were seeded on prepared collagen hydrogels at a density of 1000 crypts $well^{-1}$ in a standard 6-well plate. Collagen hydrogels for expansion were prepared with rat tail, Collagen Type 1 (354236, Corning), diluted to 1 mg $mL^{-1}$ in neutralization buffer. The final solution concentrations were: 1× PBS, 20 mM HEPES (25-060-CI, Corning), 53 mM Sodium Bicarbonate (25-035-CI, Corning), and NaOH equimolar to acetic acid, at pH 7.4. 1 mL of neutralized collagen was added to each well of a standard 6 well plate and set at 37 °C and 5% $CO_2$ for 1 h. The hydrogels were stored in sterile PBS at room temperature until use.

During expansion, cells were grown in maintenance medium (Table S1), which was changed every 48 hours. When cell confluency reached ~80% the monolayers were passaged for further expansion or use in experiments. Cells were passaged according to previously reported protocols [1, 2]. Briefly, hydrogels were digested with collagenase Type IV (500 U $mL^{-1}$, LS004189, Worthington Biochemical Corp.), followed by rinsing of the cells with 1× PBS and treatment with 0.5 mM EDTA and 10 µM Y-27632 in 1× PBS to break the large segments of monolayers into smaller pieces before re-plating at a surface area ratio of 1:3 for expansion. Cells were used for experiments between P5 and P15.

*Live/Dead Imaging Assay*

After 4 d of magnetic hydrogel culture, the magnetic agarose disks (if present) were removed, and the cells were rinsed once with 1× PBS before assaying for live/dead cellular markers. 2 mL of a staining solution consisting of 2 µM calcein-AM (354217, Corning), 2 µg $mL^{-1}$ propidium iodide (P3566, Life Technologies), and 2 µg $mL^{-1}$ Hoechst 33342 (B2261, Millipore-Sigma) in 1× PBS was added to the luminal compartment and incubated for 20 min in a humidified $CO_2$ incubator at 37 °C. The solution was aspirated, and the cells were overlaid with 1× PBS. The entire insert was placed on a glass coverslip for immediate imaging from the basal face of the insert membrane.

*Immunofluorescence Labeling*

To assess cellular lineage allocation, cultures were stained for alkaline phosphatase (ALP) activity and Muc2 presence. Directly after magnetic hydrogel culture, the cells were rinsed with 1× PBS and incubated with red ALP substrate (SK-5100, Vector Laboratories) in 150 mM Trizma buffer in the luminal reservoir for 30 min at 37 °C. Then cells were rinsed with 1× PBS, fixed in 4 % paraformaldehyde for 15 min, permeabilized with 0.5% (v/v) Triton X-100 in PBS for 20 min, and rinsed three times with a 0.75% (w/v) glycine/PBS solution. Samples were blocked with 10% donkey serum (017-000-021, Jackson Immunoresearch) in IF buffer for 90 min at room temperature. AlexaFluor 488-conjugated Muc2 antibody (1:200, sc-515032 AF488, Santa Cruz Biotechnology, Inc.) and 2 µg $mL^{-1}$ Hoechst 33342 (B2261, Millipore-Sigma) were simultaneously incubated in the 10% serum solution in the luminal reservoir 16 h at 4 °C. After a final rinse with 1× PBS, the cultures were stored in 1× PBS containing 0.1% (w/v) sodium azide.

For F-actin labeling, the cells were fixed in 4% paraformaldehyde for 15 min, permeabilized with 0.5% (v/v) Triton X-100 in PBS for 20 min and rinsed three times with 0.75% (w/v) glycine/PBS. The surfaces were blocked with 10% (v/v) donkey serum in IF buffer for 90 min at room temperature. Nuclei were stained with 2 µg $mL^{-1}$ Hoechst 33342 for 20 min at room temperature, and actin was labeled using ActinGreen 488 following the manufacturer-provided protocol (R37110, ThermoFisher Scientific). After a final rinse with PBS, the cultures were stored in PBS containing 0.1% (w/v) sodium azide.

For E-cadherin and zonula occludens-1 (ZO-1) labeling, the cultures were rinsed with PBS and fixed in ice-cold methanol at -20 °C for 1 h. All surfaces were blocked with 10% (v/v) donkey serum in IF buffer for 90 min at room temperature, prior to primary antibody incubation for 16 h at 4 °C. The primary antibodies used were: ZO-1 rabbit pAb (1:1000, 21773-1-AP, Proteintech Group, Inc.) and E-cadherin rabbit pAb (1:200, sc-7870, Santa Cruz Biotechnology, Inc.). Thereafter, the cells were rinsed with PBS and subjected to secondary antibody staining with AlexaFluor 594 donkey anti-rabbit antibody (1:500, R37119, ThermoFisher Scientific) for 45 min at room temperature. Nuclei were stained with 2 µg $mL^{-1}$ Hoechst 33342 for 1 h at room temperature, and after a final rinse with PBS, the cultures were stored in PBS containing 0.1% (w/v) sodium azide.

*Nuclear Morphology Image Analysis*

Nuclear feature extraction utilized Hoechst $33342^{+}$ areas and an automated image analysis pipeline. *Z*-series optical stacks were subjected to three-dimensional constrained iterative deconvolution in cellSens

Dimension v1.18 (Olympus Corp.) utilizing 5 iterations of an advanced maximum likelihood algorithm, with automatic denoising, background removal, and edge protection options enabled. The deconvolved images were flattened into two-dimensional representations through maximum intensity *z*-projections, and thereafter imported into CellProfiler v4.0.3 for further processing and segmentation [3]. Each two-dimensional image was flat-field corrected using a gaussian-smoothed illumination correction function (150 px filter size). The corrected images were converted to binary and segmented with the "Identify Primary Objects" module, using global 2-class Otsu threshold selection, an approximated object diameter range of 20-100 px, and shape-based de-clumping of touching objects. The area, perimeter, circularity ("FormFactor," $4\pi \times \text{area} \times \text{perimeter}^{-2}$), solidity, major/minor axis lengths, and internuclear distances ("FirstClosestDistance") for all identified objects were measured and exported to a spreadsheet.

### TABLES

#### Table S1. Formulations of primary cell culture media

| Component | Supplier Information | Final Concentration | | |
|---|---|---|---|---|
| | | *Maintenance Medium (MM)* | *Expansion Medium (EM)* | *Differentiation Medium (DM)* |
| L-WRN conditioned medium[a] | CRL-3276, ATCC (for cell line) | 50% (v/v) | 50% (v/v) | --- |
| Advanced DMEM/F12 | 12634028, ThermoFisher Scientific | 50% (v/v) | 50% (v/v) | 100% (v/v) |
| Glutamax | 35050061, ThermoFisher Scientific | 1× | 1× | 1× |
| HEPES | 25-060-CI, Corning | 10 mM | 10 mM | 10 mM |
| Primocin | ant-pm-2, Invivogen | 50 µg $mL^{-1}$ | 50 µg $mL^{-1}$ | 50 µg $mL^{-1}$ |
| N-acetylcysteine | 194603, MPBio | 1.25 mM | 1.25 mM | 1.25 mM |
| Epidermal growth factor, murine | 315-09, Peprotech | 50 ng $mL^{-1}$ | 50 ng $mL^{-1}$ | 50 ng $mL^{-1}$ |
| Nicotinamide | N0636, Millipore-Sigma | --- | 10 mM | --- |
| B27 | 17504044, ThermoFisher Scientific | 1× | 1× | --- |
| Gastrin | AS-64149, Anaspec | 10 nM | 10 nM | --- |
| Prostaglandin E2 | 14010, Cayman Chemical | --- | 10 nM | --- |
| A 83-01 | SML0788, Millipore-Sigma | 500 nM | --- | 500 nM |
| SB202190 | S-1700, LC Labs | 3 µM | 3 µM | --- |
| Y-27632 | A3008-200, ApexBio | 10 µM | 10 µM[b] | --- |
| FBS (Heat inactivated)[c] | S11050, Atlanta Biologicals | 10% (v/v) | 10% (v/v) | 0% (v/v) |

[a] Prepared by culture of the L-WRN cell line (CRL-3276, ATCC) and harvesting of media containing Wnt3a, r-spondin 2, and noggin. Refer to the protocol published by Stappenbeck and colleagues for details [4, 5].
[b] While Y-27632 is always present in MM, it is only included during the initial 24 h of culture in EM for prevention of dissociation-induced apoptosis. Thereafter, it is excluded for the remainder of experiments.
[c] FBS is included in the L-WRN conditioned medium at a concentration of 20% (diluted 50% during final media preparation for MM, EM, and SM). Therefore, no additional FBS is added to MM or EM. Heat inactivation is conducted in-house by incubating at 56°C for 25 min.

**Table S2. Quality control for cell morphology image analysis pipeline**

| Dataset | **Over-segmented objects** (req. join operation) | **Under-segmented objects** (req. split operation) | **Misidentified objects** (req. deletion) |
|---|---|---|---|
| Total (*n* = 60 images) | 4.0 ± 4.3% (7 ± 6 cells $image^{-1}$) | 2.0 ± 3.3% (4 ± 6 cells $image^{-1}$) | 1.1 ± 1.5% (2 ± 2 cells $image^{-1}$) |
| Static control (*n* = 30 images) | 2.1 ± 1.8% (5 ± 3 cells $image^{-1}$) | 2.1 ± 2.8% (5 ± 7 cells $image^{-1}$) | 0.6 ± 0.7% (2 ± 2 cells $image^{-1}$) |
| Magnetic agarose (*n* = 30 images) | 6.0 ± 5.1% (10 ± 7 cells $image^{-1}$) | 2.1 ± 3.8% (4 ± 6 cells $image^{-1}$) | 1.6 ± 1.9% (3 ± 3 cells $image^{-1}$) |

**Table S3. Cell morphology summary statistics**

| | **All cells (*n* = 7927 static control cells, *n* = 5789 magnetic agarose cells)** | | | **Data aggregated by technical replicate[a] (*n* = 5 Transwells $condition^{-1}$)** | | |
|---|---|---|---|---|---|---|
| ***Morphological parameter*** | ***Mean difference*** [95% CI][b] | ***Hedges' g*** [95% CI][b] | ***p*-value** (two-tailed Mann-Whitney) | ***Mean difference*** [95% CI][b] | ***Hedges' g*** [95% CI][b] | ***p*-value** (two-tailed *t*-test) |
| Area ($\mu m^2$) | 41.84 [38.43, 45.37] | 0.42 [0.39, 0.46] | $<2\times10^{-16}$ | 40.1 [3.77, 72.2] | 1.17 [-0.113, 2.8] | 0.074 |
| Perimeter (µm) | 9.17 [8.50, 9.87] | 0.47 [0.44, 0.51] | $<2\times10^{-16}$ | 8.82 [1.45, 15.9] | 1.22 [0.053, 2.9] | 0.065 |
| Major axis length (µm) | 3.78 [3.5, 4.06] | 0.48 [0.43, 0.52] | $<2\times10^{-16}$ | 3.68 [0.981, 6.37] | 1.37 [0.257, 2.9] | 0.043 |
| Minor axis length (µm) | 0.903 [0.781, 1.03] | 0.253 [0.218, 0.288] | $<2\times10^{-16}$ | 0.809 [-0.323, 1.7] | 0.801 [-0.568, 2.66] | 0.2 |
| Circularity | -0.041 [-0.045, -0.037] | -0.34 [-0.30, -0.37] | $<2\times10^{-16}$ | -0.0398 [-0.0646, -0.0124] | -1.5 [-3.09, -0.15] | 0.031 |
| Compactness | 0.111 [0.0446, 0.133] | 0.0927 [0.0217, 0.292] | $<2\times10^{-16}$ | 0.115 [0.0517, 0.185] | 1.72 [0.68, 2.8] | 0.017 |

[a] Cell morphology data was aggregated by technical replicate according to the SuperPlots method described by Lord S.J., et al [6]. The ggforce, ggbeeswarm, ggpubr, and Hmisc packages in RStudio were used for SuperPlot generation and significance testing.
[b] Absolute and relative effect sizes with 95% confidence intervals were calculated by the methods of Ho J., et al. using the dabest v0.3.0 package in RStudio with 20,000 bootstrap repetitions [7].

**Table S4. Nuclear morphology summary statistics**

| | **All nuclei (*n* = 8709 static control nuclei, *n* = 6616 magnetic agarose nuclei)** | | | **Data aggregated by technical replicate[a] (*n* = 5 Transwells condition$^{-1}$)** | | |
|---|---|---|---|---|---|---|
| ***Morphological parameter*** | ***Mean difference*** [95% CI][b] | ***Hedges' g*** [95% CI][b] | ***p*-value** (two-tailed Mann-Whitney) | ***Mean difference*** [95% CI][b] | ***Hedges' g*** [95% CI][b] | ***p*-value** (two-tailed *t*-test) |
| Area (μm$^2$) | 9.21 [8.58, 9.85] | 0.48 [0.45, 0.51] | <2×10$^{-16}$ | 8.88 [1.84, 17.3] | 1.16 [-0.0387, 2.41] | 0.077 |
| Perimeter (μm) | 2.45 [2.29, 2.62] | 0.5 [0.467, 0.534] | <2×10$^{-16}$ | 2.38 [0.602, 4.49] | 1.23 [0.0979, 2.49] | 0.063 |
| Major axis length (μm) | 1.11 [1.04, 1.18] | 0.523 [0.491, 0.557] | <2×10$^{-16}$ | 1.09 [0.31, 1.97] | 1.31 [0.183, 2.6] | 0.051 |
| Minor axis length (μm) | 0.275 [0.239, 0.311] | 0.246 [0.214, 0.278] | <2×10$^{-16}$ | 0.251 [-0.0456, 0.587] | 0.787 [-0.602, 2.06] | 0.21 |
| Circularity | -0.014 [-0.017, -0.012] | -0.18 [-0.21, -0.15] | <2×10$^{-16}$ | -0.017 [-0.0283, -0.00611] | -1.53 [-2.93, -0.263] | 0.031 |
| Compactness | 0.0247 [0.0188, 0.0307] | 0.134 [0.101, 0.166] | <2×10$^{-16}$ | 0.0261 [0.00885, 0.0462] | 1.41 [0.195, 2.63] | 0.042 |
| Internuclear distance (μm) | 0.95 [0.88, 1.03] | 0.40 [0.37, 0.44] | <2×10$^{-16}$ | 0.89 [-0.0956, 1.75] | 0.959 [-0.353, 2.65] | 0.13 |

[a] Cell morphology data was aggregated by technical replicate according to the SuperPlots method described by Lord S.J., et al [6]. The ggforce, ggbeeswarm, ggpubr, and Hmisc packages in RStudio were used for SuperPlot generation and significance testing.
[b] Absolute and relative effect sizes with 95% confidence intervals were calculated by the methods of Ho J., et al. using the dabest v0.3.0 package in RStudio with 20,000 bootstrap repetitions [7].

### FIGURES

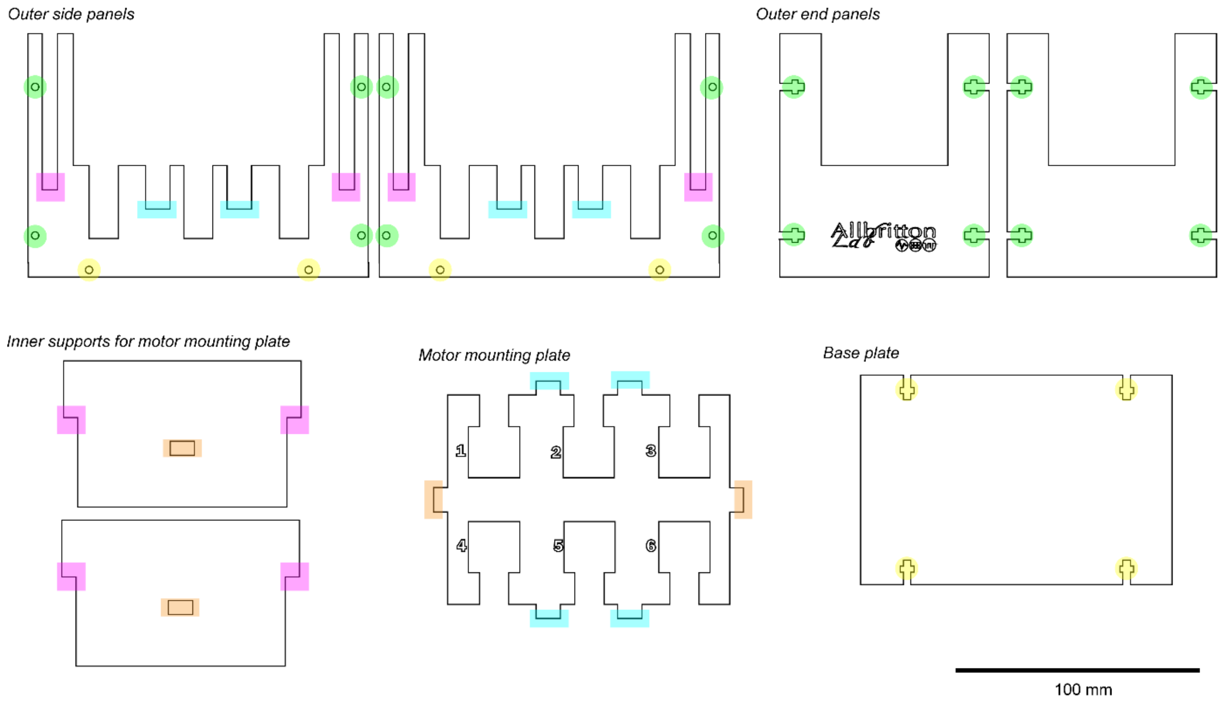


**Figure S1.** *Acrylic frame schematics.* The above two-dimensional panels were laser cut from clear cast acrylic and assembled with stainless steel socket cap screws. The respective joining regions are color coded for clarity of assembly.

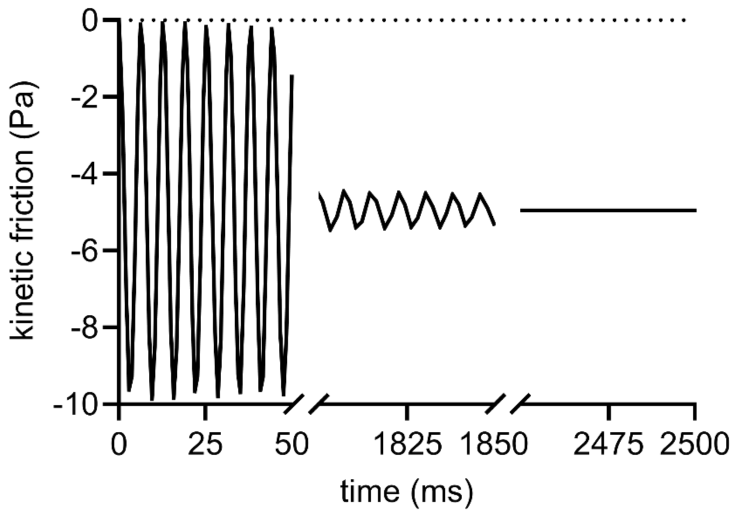


**Figure S2.** *Dependence of average kinetic friction on time within COMSOL model.* By 2500 ms, the frictional force converges to a constant value from which all data in the manuscript (*i.e.*, shear stress plots and average kinetic friction values) are reported.

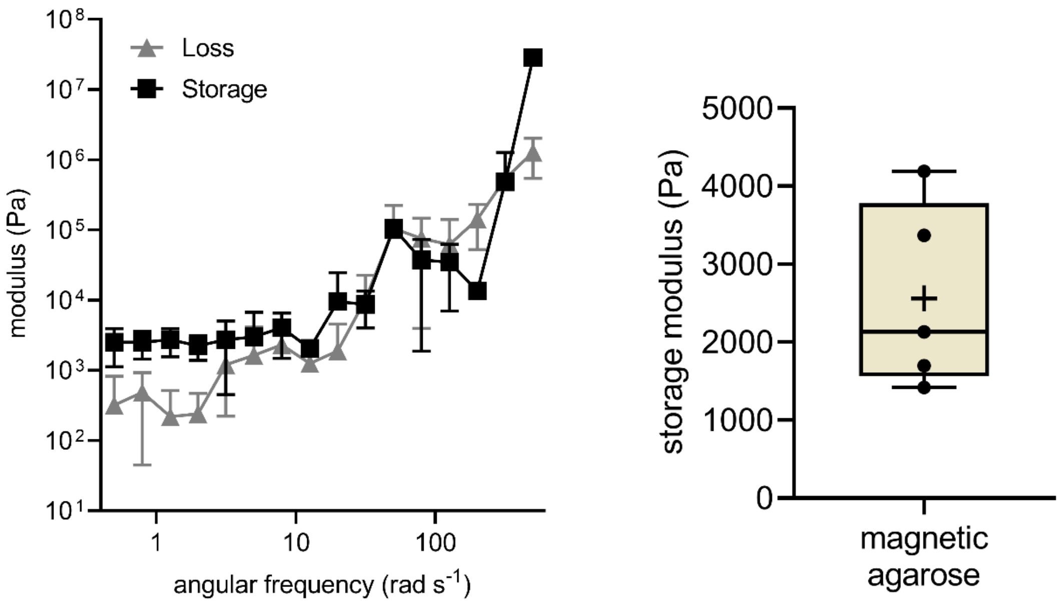


**Figure S3.** Rheological analysis of the magnetic agarose (1.7% w/v agarose, 10 mg $mL^{-1}$ magnetic nanoparticles) was conducted on an Anton Paar MCR-301 rheometer with an 8 mm diameter parallel plate with a 0.5 mm gap size. Frequency sweeps were collected between 0.5 and 500 rad $s^{-1}$ with 0.01% strain amplitude. Storage moduli (G' = 2562 ± 1177 Pa, mean ± S.D., *n* = 5 hydrogels) were averaged across the linear viscoelastic region of the frequency sweep, which ranged from 0.5-2 rad $s^{-1}$. Assuming a Poisson's Ratio of 0.49 [8], the mean storage modulus can be converted by the Theory of Elasticity to an approximate Young's modulus of 7635 ± 3507 Pa. The box plot crossbars represent the sample median, 25th percentile, and 75th percentile, with the whiskers depicting the minimum and maximum measurements, and "+" depicting the sample mean.

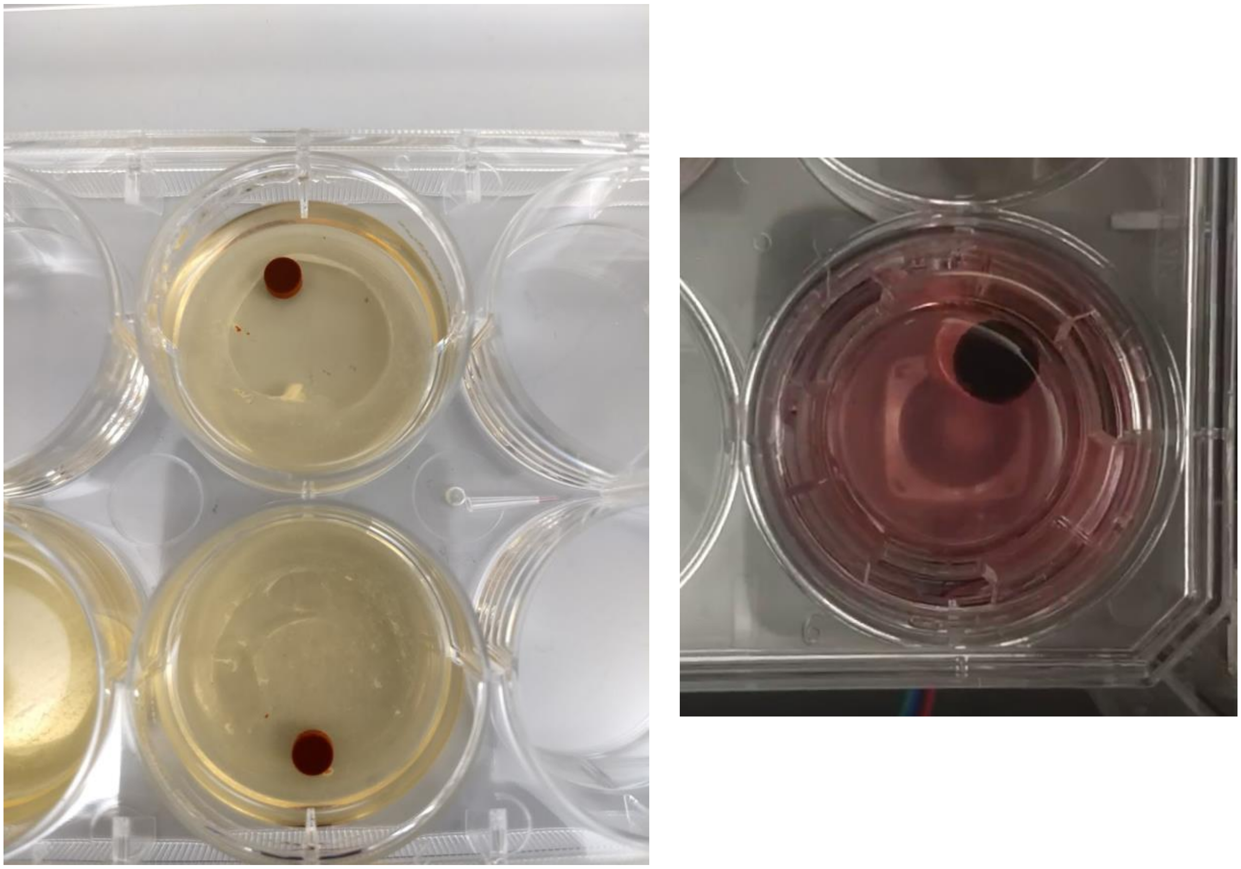

**Figure S4.** *Deformation of neutralized collagen hydrogels by magnetic agarose.* (Left) After rotational exposure of the hydrogels to the magnetic agarose, the neutralized collagen hydrogel is permanently etched due to its much lower stiffness (elastic modulus <100 Pa). These results necessitate the use of a stronger support for the intestinal epithelial cells in this platform. (Right) Conversely, porous membranes with a thin surface coating of collagen are not etched or deformed by magnetic agarose rotation. These surface coatings retain the ability to support intestinal cell attachment and expansion.

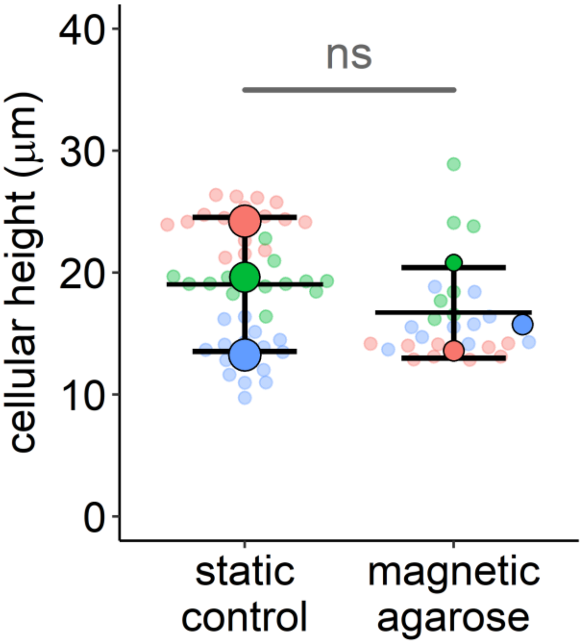


**Figure S5.** *Comparison of cellular heights.* Heights were measured using Fiji [9] from *xz*-slices of three replicate fluorescent image stacks in which F-actin and DNA were probed. The average height of the static control cells was 19.0 ± 4.9 µm ($n$ = 47 cells) and the average height of the magnetic agarose exposed cells was 16.4 ± 3.9 µm ($n$ = 28 cells). These results were aggregated by technical replicate and compared using an unpaired, two-tailed t-test ($p$ = 0.58) in RStudio [6]. The transparent data points represent individual cells and are color-coded by technical replicate, while the solid data points represent the mean value for each technical replicate and are sized by the number cells identified in each image.
